# Novel extracellular matrix architecture on excitatory neurons revealed by HaloTag-HAPLN1

**DOI:** 10.1101/2024.03.29.587384

**Authors:** Igal Sterin, Ava Niazi, Jennifer Kim, Joosang Park, Sungjin Park

**Author notes:** **Correspondence** Sungjin Park, Department of Neurobiology, University of Utah School of Medicine, Salt Lake City, UT 84112.

## Abstract

The brain’s extracellular matrix (ECM) regulates neuronal plasticity and animal behavior. ECM staining shows an aggregated pattern in a net-like structure around a subset of neurons and diffuse staining in the interstitial matrix. However, understanding the structural features of ECM deposition across various neuronal types and subcellular compartments remains limited. To visualize the organization pattern and assembly process of the hyaluronan-scaffolded ECM in the brain, we fused a HaloTag to HAPLN1, which links hyaluronan and proteoglycans. Expression or application of the probe enables us to identify spatial and temporal regulation of ECM deposition and heterogeneity in ECM aggregation among neuronal populations. Dual-color birthdating shows the ECM assembly process in culture and *in vivo.* Sparse expression in vivo reveals novel forms of ECM architecture around excitatory neurons and developmentally regulated dendritic ECM. Overall, our study uncovers extensive structural features of the brain’ ECM, suggesting diverse roles in regulating neuronal plasticity.

## Introduction

The brain’s extracellular matrix (ECM) is implicated in numerous processes from providing a niche for neurogenesis to regulating the integrity of the blood-brain barrier [1-3]. Notably, the ECM regulates plasticity and behavior [4-7]. Digestion of the main sugar modification on the major protein subunits of the ECM, chondroitin sulfate on chondroitin sulfate proteoglycans (CSPGs), with chondroitinase, restores juvenile levels of plasticity in a variety of paradigms, including ocular dominance plasticity, auditory learning, and fear memory consolidation [8-11]. Also, aged animals and animals in neurodegenerative models display increased memory performance after digestion of the ECM, overexpression of chondroitin-6-transferase, or ECM component knockouts [6, 7, 12-14]. Since loosening the ECM through pharmacological, genetic, or behavioral manipulations leads to increases in plasticity, the ECM is considered a general barrier to neuronal plasticity [15-21]. The ECM in the brain parenchyma is scaffolded on up to mega-Dalton polymers of hyaluronic acid (HA), a sugar molecule that is secreted from cells by transmembrane hyaluronan synthase [16, 22-24]. HA forms a large complex with the lectican family of CSPGs, including aggrecan, brevican, versican, and neurocan, which are crosslinked by the glycoprotein tenascin-r and stabilized by the hyaluronan and proteoglycan link (HAPLN) family [2, 14, 25-28]. This HA-based ECM is intrinsically flexible and makes ultrastructural identification of the ECM difficult. Staining against CSPGs or chondroitin sulfate 5,6 (CS-56) shows diffuse staining in most of the brain parenchyma and net-like aggregation around certain cells [23, 29-33]. Conversely, staining with *wisteria floribunda agglutinin* (WFA), a plant lectin that binds to aggrecan glycosaminoglycans (GAGs), displays highly contrasting, dense net-like coats of aggregated ECM termed perineuronal nets (PNN) with little staining of the interstitial matrix [34].

WFA (+) PNNs are predominantly found around the soma and proximal neurites of parvalbumin (PV) interneurons and a subset of excitatory neurons in specific brain regions, including the hippocampal CA2 and amygdala [5, 9, 34]. Since both chondroitinase digestion and genetic knockout of ECM components reduce WFA (+) labeling and alter the electrophysiological properties of PV neurons [19, 35, 36], WFA (+) PNNs around PV neurons are considered the functional units responsible for the ECM’s ability to regulate plasticity. This has led to a model where PNNs grow denser around PV neurons, decreasing the number or strength of the upstream inhibitory contacts on their soma. Digestion of WFA (+) PNN around PV neurons leads to an increase in the inhibitory input to the PV neurons and subsequent disinhibition of downstream excitatory neurons [3, 16, 17, 19, 36-38]. However, since chondroitinase digestion and global knockout of ECM components impact not only WFA (+) PNN but also the entire matrix, the specific role of PV-PNNs in neuronal plasticity and behavior remains to be determined.

Notably, antibodies against other CSPGs and CS GAGs show distinct clustering patterns that partially overlap with WFA (+) PNNs [23, 29-33], suggesting that there are other types of structured ECMs that cannot be stained with WFA. Moreover, PNN-like clusters could be more prevalent since the identification of the contrasting clusters by global staining of CSPGs or CS GAGs can be obscured in brain areas where neurons are densely packed such as the hippocampal pyramidal cell body layers. While the interstitial matrix is considered a mostly unstructured gel, detailed examination has remained challenging. Recent studies show that ECM’s role is not restricted to the neuronal soma. It regulates synaptic plasticity by regulating neurotransmitters’ diffusibility in and beyond the synaptic cleft [39-41], AMPAR receptor mobility [42], and physiological remodeling of the ECM in the hippocampus can increase plasticity [43].

To extensively identify ECM architecture, we developed a novel genetic tool that labels the brain’s HA-based ECM. We chose HAPLN1, a link protein which binds the HA backbone of the ECM and CSPGs [31, 44, 45] and implicated in ECM plasticity phenotypes: HAPLN1 knockout animals have improved memory performance in advanced age and attenuated WFA (+) PNNs [6, 16, 27]. Fusion of HaloTag [46, 47] to HAPLN1 allows us to visualize the ECM in culture and *in vivo*. We provide evidence for spatial and temporal regulation of ECM deposition that is heterogeneous between individual neurons in culture and *in vivo.* We observed a novel ECM architecture around pyramidal neurons of the hippocampus that did not overlap with WFA (+) PNNs as well as aggregation of the ECM in the interstitial matrix that was developmentally regulated. Based on our observations, we posit a model wherein the ECM regulates the balance between flexibility and stability of synapses on both excitatory and inhibitory neurons.

## Results

### HALO-Tagged HAPLN1, H-Link, can be visualized in live neurons and reveals maturation of neuronal ECM

Staining of cultured neurons with conventional ECM markers shows minimal organized signal till days in vitro (DIV) 21 [15, 22, 48, 49]. This could be due to the delay of ECM deposition in culture, or inadequate visualization of ECM assembly. To differentiate these possibilities, we aimed to trace the localization of a link protein that stabilizes the interaction between HA and CSPGs. We inserted a HaloTag between the signal peptide and the mature peptide of HAPLN1 (hyaluronan proteoglycan link protein) downstream of a GFP-T2A sequence to label infected neurons (Figure 1A). We reasoned that a tagged version would compete with the endogenous link proteins for binding sites between the HA and CSPGs, thus revealing ECM structure. We term this fusion protein H-Link, for Halo-Link protein.

**Figure 1:**
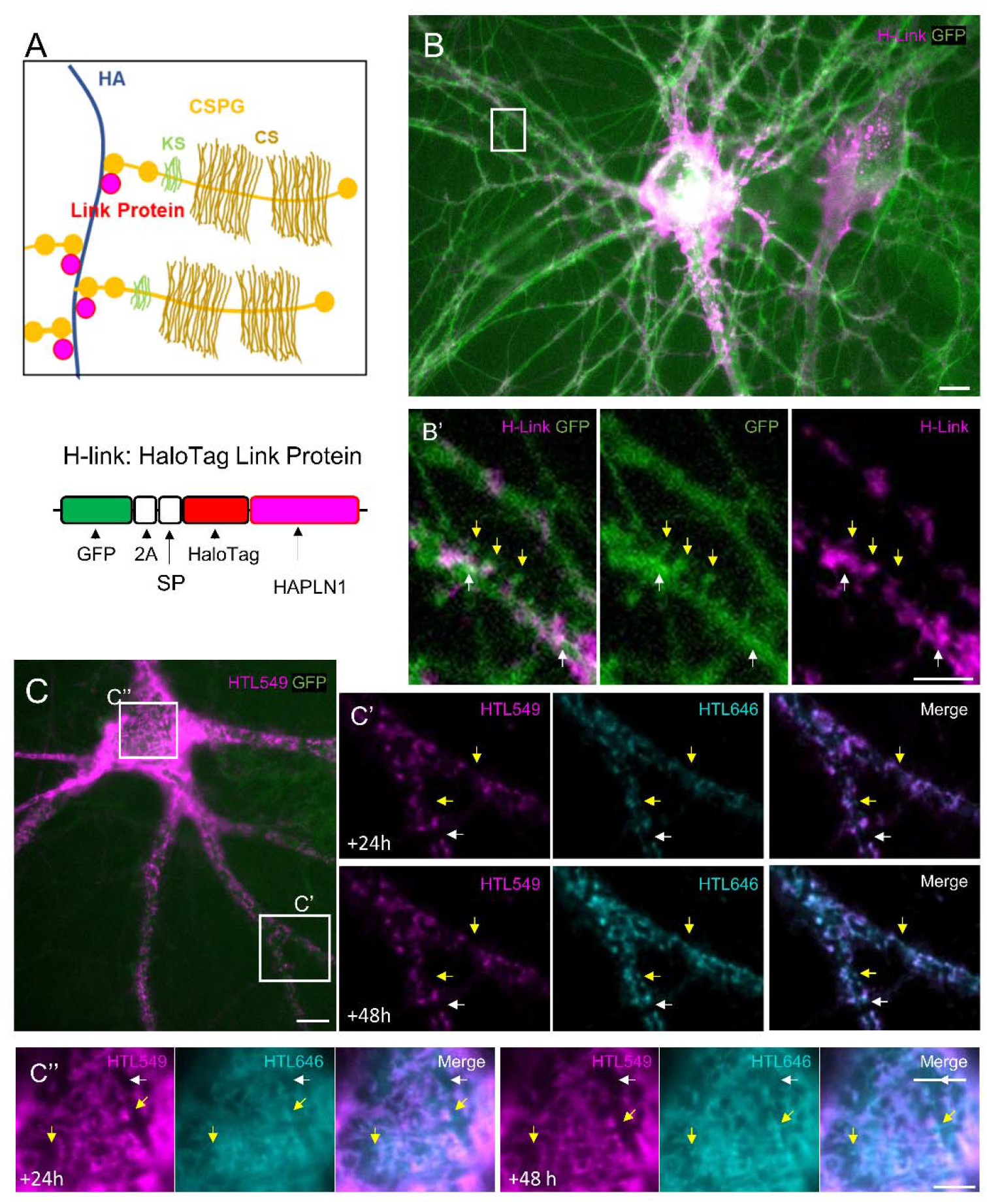
HaloTag HAPLN1, H-Link, can be visualized in live neurons and reveals maturation of the ECM. **(A)** *Top*: simplified diagram of the extracellular space with the hyaluronic acid (HA) backbone of the ECM bound by chondroitin sulfate proteoglycans (CSPGs) with keratin sulphate (KS) and chondroitin sulfate (CS) posttranslational modifications. Smaller link proteins bind both CSPGs and HA. *Bottom:* Schematic of novel genetic probe for visualizing the ECM consisting of a HaloTag inserted after the endogenous signal peptide (SP) of hyaluronan proteoglycan link protein 1 (HAPLN1) as well as a GFP and T2A sequence as an expression marker. **(B)** Primary rat hippocampal cultures were transduced with AAV H-Link on DIV3, HTL549 was added on DIV9, and imaging on live neurons began on DIV10. Representative image of a DIV10 neuron showing GFP expression visible in green with HTL549 signal showing dense H-Link aggregation around the soma and proximal dendrites. Scale bar 50 µm. **(B’)** Inset of B, with H-Link signal occluded from dendritic spine heads (yellow arrows) and aggregating around neurite crossovers (white arrows). Scale bar 5 µm. **(C)** Representative image of DIV8 neuron transduced with H-Link AAV on DIV1 and HTL549 added on DIV7. Net-like pattern of H-link is visible in the neuronal soma, with punctate signal along proximal dendrites. Scale bar 10 µm. **(C’, C’’)** Zoom in on process (**C’**) or soma (**C’’**) in **C** at 24 or 48 hours after HTL646 addition on DIV9 showing areas where there is HTL646 but no HTL549 (yellow arrows) and areas where there is neither HTL (white arrows) Scale bar 5 µm.

Rat hippocampal cultures expressed GFP visibly by DIV7 after being transduced with AAV H-Link on DIV1-3 (Data not shown). H-Link visualized in live cells by adding HaloTag ligand (HTL) conjugated to Janelia fluorophore 549 (HTL549) (Figure 1B) is incorporated into neuronal ECM overnight. H-Link signal was denser around the soma and proximal neurites and punctated at distal neurites. Upon closer examination, we observed that the signal was absent from dendritic spine heads. Instead, there was an aggregation of HTL signal at crossover sites of neurites (Figure 1B’). H-Link signal was stable over several days (Figure 1C-C’’). Taking advantage of the versatility of the HaloTag, we birthdated the overexpressed H-Link by dual HTL labeling (Movie 1). Adding HTL conjugated to Janelia fluorophore 646 (HTL646) 48 hours after HTL549 to transduced neurons resulted in overlapping and separated signal, as well as areas that lacked either signal, showing the maturation process of ECM deposition (Figure 1C-C’’ arrows).

### The application of exogenous H-Link reveals heterogeneity in ECM aggregation in cultured neurons

Since there is no established method to interrogate the ECM of early cultures, we could not exclude the possibility that overexpression of H-Link impacts ECM development. To circumvent overexpression, we overexpressed H-link in HEK293T cells and transferred the conditioned media (CM) to more mature neurons, thus H-Link would bind to existing ECM (Figure 2). We preincubated the CM with HTL549 and applied it to neurons on DIV13. Neurons incubated overnight with exogenous H-link show staining similar to the overexpressed H-link: dense soma aggregation and punctate along distal neurites (Figure 2A-B). Neurons incubated with the CM of untransfected HEK293T cells or HTL549 alone showed no staining pattern (Figure S1A-C, J). Cultures pretreated with hyaluronidase, a HA degrading enzyme, had a clear decrease in signal after overnight incubation with H-Link CM (Figure S1D, J). To further determine the specificity of the binding of H-Link to HA, we created two mutants of H-link: an alanine mutant where the most conserved six amino acids in 2nd HA binding proteins were mutated to alanine and a deletion mutant with a premature stop codon, truncating the C-terminus by 68 amino acids of the HA binding domain (Figure S1E) [44, 45, 50]. H-Link mutants did not display clustered patterns (not shown). Western blots validated that the mutants were expressed and secreted (Figure S1I), although to a lesser degree than the unmutated H-Link. To rule out the possibility that the absence of signal from H-Link mutants was due to the reduced section from HEK293T cells, we added five times more H-Link mutant CM but did not observe the clustered pattern around neurons (Figure S1F-H, J). Overall, these observations indicate that diffusible H-Link is incorporated into existing neuronal ECM clusters through its ability to bind intact HA chains.

**Figure 2:**
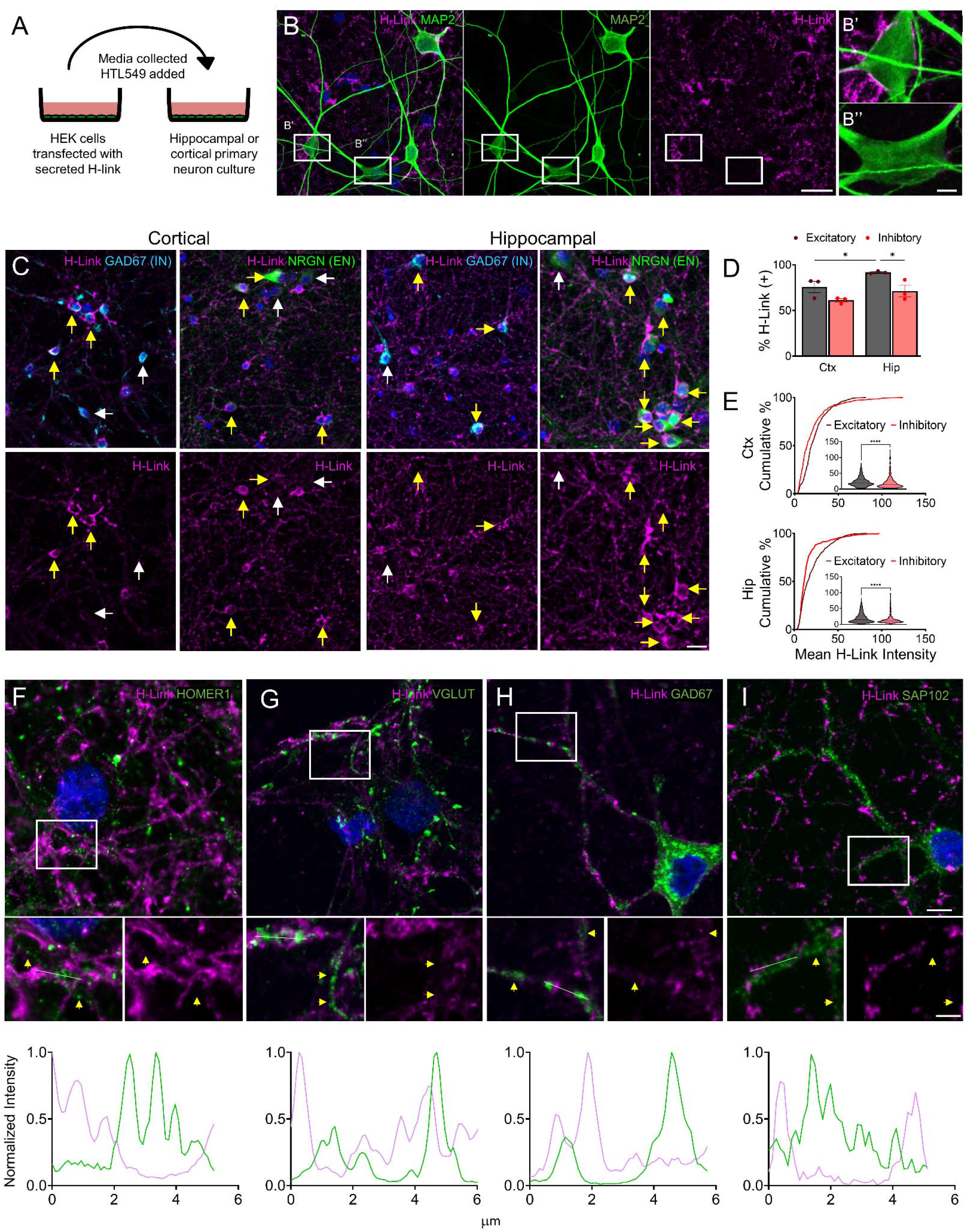
Exogenous H-Link probe reveals heterogeneity in ECM aggregation in cultured neurons. **(A)** Schematic of H-Link as an exogenous probe wherein H-Link is expressed in HEK293T cells, conditioned media (CM) from these cells is collected and incubated in HTL549 before being added to primary rodent neuronal cultures. **(B)** Representative max projection of DIV14 rat hippocampal cultures incubated overnight in H-Link CM shows heterogeneous H-Link aggregation between neuronal somas and diffuse or punctate staining along distal dendrites. Magenta: HTL549 (H-Link); Green: MAP2. Scale bar 25 µm. **(B’ B’’)** inset from **B** highlighting the heterogeneity of soma coverage: **B’** has deposition around the soma, and **B’’** does not. Scale bar 5 µm. **(C)** Representative max projections from DIV14-18 cortical (*Left*) and hippocampal (*Right*) cultures stained for excitatory neurons (EN) with neurogranin (NRGN), or inhibitory neurons (IN) with GAD67 after overnight incubation in H-Link CM. *Top*: merged image with DAPI *Bottom:* paired H-Link only. Yellow arrows denote H-Link positive neurons whereas white arrows denote H-Link negative neurons. Scale bar 10 µm. **(D)** Quantification of % of H-Link (+) neurons among different neuronal populations. Hippocampal excitatory neurons showed a significantly higher % of H-Link (+) neurons compared to other populations. cortical excitatory (CE), 75.74 ± 6.2%; cortical inhibitory (CI) 61.12 ± 1.822%; hippocampal excitatory (HI) 91.68 ± 0.748%; hippocampal inhibitory (HI) 71.33 ± 6.347, % ± SEM, N=3 cultures each CE=659 neurons, CI=420 neurons, HE= 2001 neurons, HI =396 neurons. Two-way ANOVA culture factor F=10.75, p=0.0305, neuron factor F= 12.08 p=0.0254. Uncorrected Fisher’s LSD: CE vs. CI p=0.1090; HE vs. HI p=0.0459; CE vs. HE p=0.0379; CI vs. HI p=0.1505. **(E)** Cumulative distribution of the mean H-Link intensity of each neuron for cortical (Ctx) (*Top*) and hippocampal (Hip) (*Bottom*) with violin plot with median and quartiles. N=3 cultures each, 10 coverslips (386 excitatory neurons and 220 inhibitory neurons) in the cortical cultures, 9 coverslips (534 excitatory neurons and 326 inhibitory neurons) in the hippocampal culture. Kolmogrov-Smirnov test, CE vs CI p<0.0001, HE vs HI p<0.0001. **(F-I)** H-Link does not localize with synaptic markers. *Top*: representative images H-Link (magenta), DAPI (blue), and indicated synaptic marker (green). *Middle*: zoomed-in view of insert. *Bottom*: line scan of H-Link (magenta) and a synaptic marker (green) of the white line in the middle image. Line scans show alternating peaks and troughs, indicating the separation of the signal. Yellow arrows show clusters of synaptic markers. Scale bar *Top* 5 µm, *Middle* 2.5 µm.

Notably, we observed clear heterogeneity of H-Link clustering among neurons, ranging from dense heavy signal to no signal at all (Figure 2B-B’’). To explore this further, we compared the populations and intensity of H-link around excitatory neurons vs inhibitory neurons in hippocampal and cortical cultures (Figure 2C). We found that a majority of the excitatory neurons in hippocampal cultures displayed H-Link clusters, whereas cortical excitatory neurons and cortical and hippocampal inhibitory neurons had a significantly smaller proportion of neurons displaying H-Link clusters (Figure 2D) (cortical excitatory (CE), 75.74 ± 6.2%; cortical inhibitory (CI) 61.12 ± 1.822%; hippocampal excitatory (HI) 91.68 ± 0.748%; hippocampal inhibitory (HI) 71.33 ± 6.347, % ± SEM, Two-way ANOVA culture factor F=10.75, p=0.0305, neuron factor F=12.08 p=0.0254. N=3 cultures, Uncorrected Fisher’s least significant difference (LSD): CE vs. CI p=0.1090; HE vs. HI p=0.0459; CE vs. HE p=0.0379; CI vs. HI p=0.1505). This observation indicates regional differences in excitatory neuronal populations capable of ECM deposition. Next, we quantified the mean intensity of H-Link deposition on individual neurons and compared the cumulative distribution within cultures. We saw that in both culture conditions, the excitatory distribution was shifted to the right, indicating a higher level of H-Link deposition than inhibitory neurons (Figure 2E) (Kolmogrov-Smirnov test, CE vs CI p<0.0001, HE vs HI p<0.0001) Interestingly, there is an inhibitory neuronal population specialized in ECM deposition, visible as a crossover and rightward shift in the cumulative plots (Figure 2E) H-link punctate signal along the neurite processes begs the question of whether this signal is co-localized with synaptic markers. H-link signal was segregated from several common synaptic markers such as inhibitory GAD67, and VGLUT, HOMER1 and SAP102 for excitatory synapses [51-54] (Figure 2F-I). Clearly, the application of H-Link identifies the neurons inducing ECM deposition and subcellular localization of ECM clusters.

### ECM is organized early, and deposition is spatially and temporally regulated

To determine when neurons organize their ECM, we performed live-cell time-lapse imaging on primary hippocampal cultures. HTL-conjugated H-Link CM was added on DIV2, imaging began on DIV3, and cultures were followed till DIV14. Over the time course, we identified three patterns of neuronal H-Link clustering: H-Link negative, early organizers, and late organizers (Figure 3A, B) (Movie 2). For example, neurons 5 and 8 do not organize ECM, neurons 1 and 6 began to organize their ECM between DIV5 and 7, and the rest of the identified neurons are late organizers, as they begin to organize their ECM after DIV7. Of note is that the probe is added early and homogenously but organized in a spatially and temporally regulated way, with positive and negative neurons, indicating that the absence of H-Link clusters in some neurons is not due to the depletion of the probe. Next, we tested whether the maintenance of H-Link clusters requires intact HA chains. We added hyaluronidase on DIV7 and saw a substantial loss of signal around early organizers (Figure 3C, n=20 neurons, paired t-test, p<0.0001). Timelapse imaging also showed H-Link aggregation was induced after neurite crossover (Figure 3D). We can watch as a growth cone crosses another neurite and, as early as two hours post-crossing, there was an aggregation of the signal at the crossover that lasted through a 24 or 48-hour imaging session (Movie 3, 4). Due to the elaborated neurite crossovers during the period when neurons begin assembling ECM clusters (DIV5 and later), the observation of new cluster formation after a new crossover was obscured. However, the contact-mediated induction suggests that ECM deposition is temporally and spatially regulated on dendrites. These observations highlight that live-cell imaging of H-Link reveals the dynamic organizational process of neuronal ECM.

**Figure 3:**
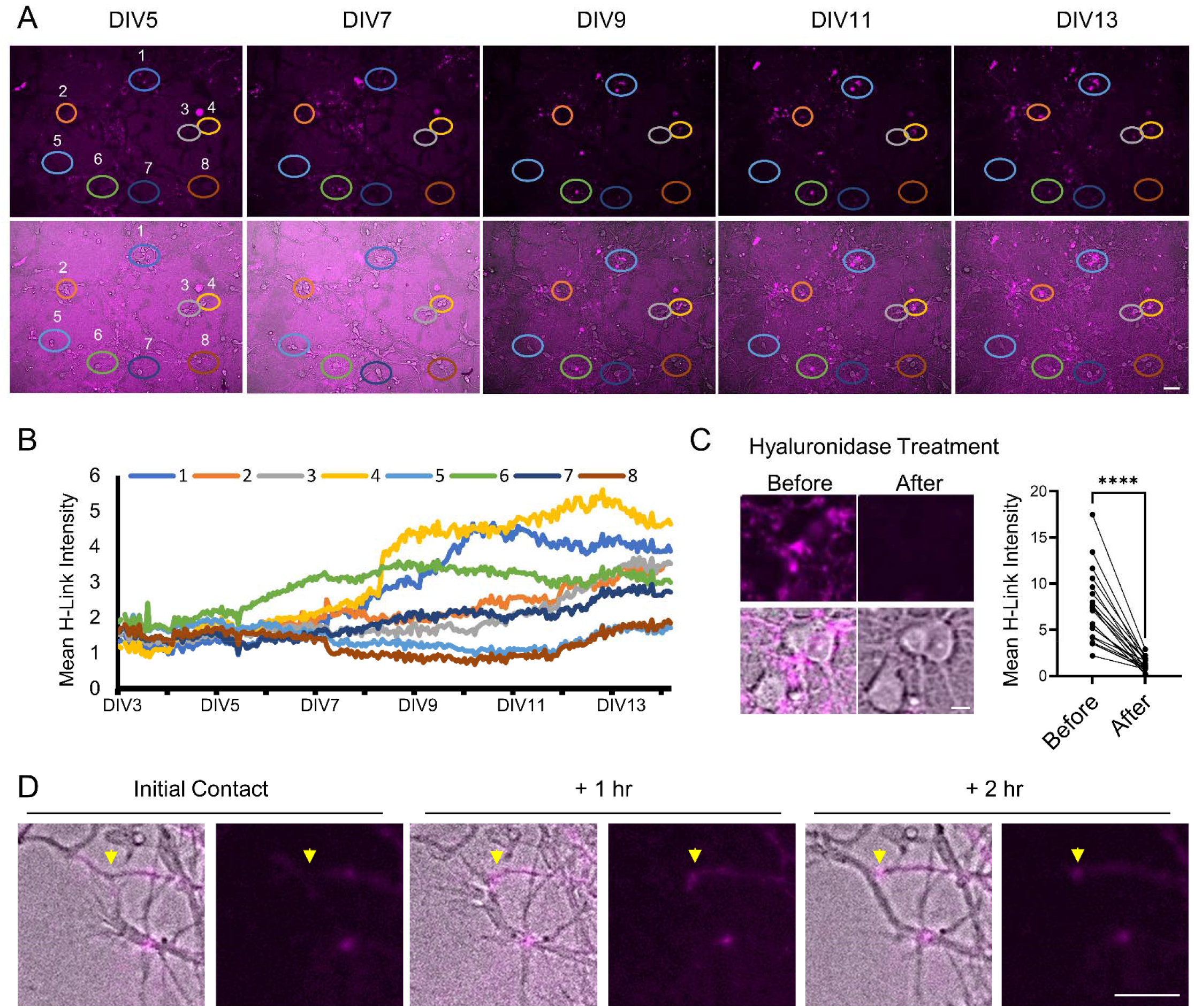
The ECM is organized early, and deposition is spatially and temporally regulated. **(A and B)** Rat hippocampal cultures were treated with H-Link CM on DIV2, and time-lapse imaging was taken (magenta: H-Link; gray: bright field) from DIV3 till DIV14 every hour. **(A)** Representative field of view from DIV5 till DIV13. Neurons are circled at each time point highlighting neuronal diversity in H-Link aggregation. Scale bar 50 µm. **(B)** Quantification of the mean fluorescence intensity of H-Link on neuronal soma shows distinct temporal patterns of H-Link aggregation on different neurons. **(C)** *Left*: Treatment of hyaluronidase (100 U/mL) caused loss of H-Link signal. Scale bar 5 µm. *Right* Quantification of 20 neurons before treatment and 7 hours after treatment shows a dramatic and significant loss of signal (paired Student’s t-test, p<0.0001). **(D)** Aggregation of H-Link follows neurite contact, where yellow arrows indicate the contact site and panels show initial contact to 2 hours later in a DIV6 culture with H-Link CM added on DIV5. Scale bar 10 µm. (**A, C,D)** Magenta: H-Link (HTL549); Gray: brightfield.

### H-Link expression *in vivo* reveals conventional and unconventional ECM

Next, we tested whether H-Link reveals ECM organization patterns *in vivo*. The promoter, infection marker, and tagging sequences are modular in the AAV vector, so we packaged AAVs that tested multiple promoters and either membrane or cytoplasmic GFP (mGFP or eGFP) with H-Link and injected them into post-natal day 1 (P1) mouse brains and examined the expression at 3 weeks. We found that HTL could label H-Link in PFA-fixed brain tissue and observed H-Link signal with aggregation around neurons that was reminiscent of WFA (+) PNN structures (Figure S2B). Conversely, staining of uninfected brain tissue with HTL did not produce signal (Figure S2A). The CAG promoter (pCAG) drove expression in both astrocytes and neurons (Figure S2B), and pGFABC1D promoter drove expression predominantly in astrocytes and leaky expression in some neurons (not shown). As expected, the Synapsin promoter (pSYN) was selective for neurons (Figure 4, S2E). The mGFP showed that the HTL signal was outside of cells (Figure S2B). Demonstrating the modularity of the tool, we swapped the HaloTag for a GFP, making GFP-Link (G-Link) which also revealed ECM in the brain (Figure S2C).

**Figure 4:**
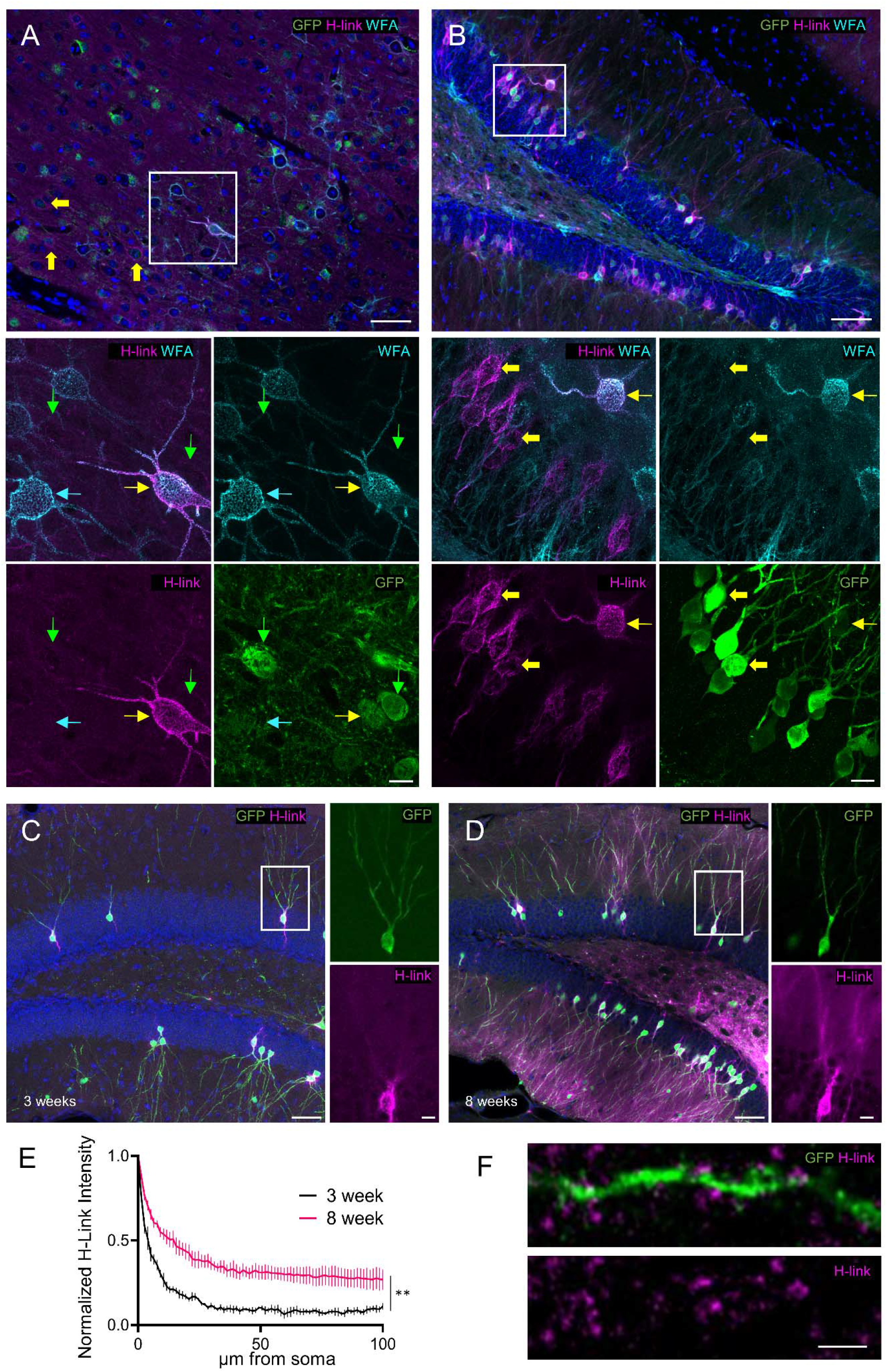
*In vivo*, H-Link reveals novel forms of ECM structure and developmental maturation on dendrites. **(A)** *Top:* Representative image of the somatosensory cortex layer IV of an 8-week animal injected with pSYN-eGFP-2A-H-Link AAV at P2. *Bottom:* Single color and WFA HTL Merge of the max projection of the inset of the *Top*. H-Link signal was detected on WFA (-) neurons (yellow block arrows) and WFA (+) neurons (yellow arrow). Green arrows indicate GFP (+), H-Link (-) neurons. Cyan arrows indicate an uninfected WFA (+) neurons. H-Link clusters on WFA (+) neurons closely overlapped with net-like WFA architecture. *Top* Scale bar 50 µm, *Bottom* Scale bar 10 µm. **(B)** *Top:* Representative image of the dentate gyrus (DG) of 8-week animal injected with pSYN-eGFP-2A-H-Link AAV at P2. *Bottom:* Single color and WFA HTL Merge max projection of inset. H-Link signal was detected on WFA (-) DG granule cells (yellow block arrows) and WFA (+) neurons (yellow arrow). H-Link clusters display conventional net-like architecture on WFA (+) neurons and a distinct unwoven fiber coat on excitatory neurons. All GFP (+) neurons have H-Link signal in the hippocampus. *Top* Scale bar 50 µm, *Bottom* Scale bar 10 µm. **(C)** Representative max projection of 3-week dentate gyrus injected with pSYN-eGFP-2A-H-Link AAV at P2 with inset showing minimal aggregation along dendrites. **(D)** Representative image of 8-week dentate gyrus injected with pSYN-eGFP-2A-H-Link AAV at P2 with inset showing increased aggregation along dendrites in the older animal. (**C, D)** Scale bar 50 µm, inset 10 µm. **(E)** Eight to ten neurons having intact dendritic branches on the cross-section and displaying bright H-Link signal were selected per animal. The corrected fluorescence intensity of the H-Link signal after background subtraction was quantified along the dendritic length and normalized to the highest intensity on the soma. Dendritic H-Link signal is significantly higher in 8 weeks compared to 3-week-old animals (N=4 animals per group, SEM error bars, Two-way repeated measure ANOVA, Distance factor F=181.5 p<0.0001; Age factor F=21.9 p=0.0034). **(F)** High-resolution Airyscan image of the DG dendrite at 8 weeks with eGFP infection marker and H-Link signal merged shows a lack of H-Link signal at spine heads with uneven distribution on the dendritic shaft and spine necks. Scale bar 2 µm.

We found that the pSYN promoter with eGFP gave the clearest signal around the soma and dendritic processes and allowed for sparse staining to distinguish expressing vs non-expressing cells. We decided to examine 8-week-old animals injected with H-Link AAV at P2 and focused on the somatosensory cortex and hippocampus for the initial characterization. Interestingly, we found that in the somatosensory cortex, there were cells that expressed GFP but showed little or no HTL staining (Figure 4A and S2E, green arrows), whereas, in the hippocampus, almost all GFP+ cells were also H-Link (+) (Figure 4B), showing heterogeneity of the ECM depending on the region. Both regions had H-Link (+) WFA (+) and H-Link (+) WFA (-) PNNs (Figure 4A, B, S2E, yellow and yellow block arrows, respectively). Upon closer examination of H-Link (+) WFA (+) neurons, the H-Link signal closely overlapped with WFA (+) PNN architecture (Figure 4A) (Movie 5). H-Link overexpression did not cause notable changes in WFA or chondroitin sulfate staining patterns compared to those on uninfected neurons (Figure 4A, yellow and cyan arrows, respectively, Figure S2D). Of note, in each brain injected, we were able to find uninfected cells that captured the diffusible H-Link (Figure S2E-G). They were predominantly observed in the cortex and were WFA (+) or WFA (-) (Figure S2E and S2G, white and pink arrows, respectively).

Interestingly, we observed a PNN-like cluster on excitatory neurons infected with H-Link. For example, in the dentate gyrus (DG) of the hippocampus, a region of low conventional PNNs labeled by WFA, we observed H-Link signal around dentate granule cells (Figure 4B). This ECM around excitatory cells displayed a distinct three-dimensional structure compared to conventional WFA (+) PNN (Movie 6), appearing more as unwoven fibers as opposed to the classical net structure. We have termed this novel form of ECM, revealed by our overexpression system, excitatory perineuronal coats (PNC). In addition to the PNC, we also noticed punctate signal extending up along the dendrites of the granule cells. (Figure 4D, F). We were interested if the puncta along the dendritic arbors of dentate granule cells matures over time, as it is known that WFA signal increases as an animal ages [3, 9]. We compared 3-week and 8-week-old animals (Figure 4C,D) and observed an increase in the coverage and intensity of the dendritic deposition normalized to the soma signal in the DG neurons (Figure 4E, Two-way repeated measure ANOVA, Distance factor F=181.5 p<0.0001; Age factor F=21.9 p=0.0034). These observations suggest that the level of ECM clustering on dendrites of excitatory neurons is developmentally regulated during the juvenile-adult transition. High-resolution Airyscan imaging of the dendritic area at 8 weeks showed that dendritic H-Link puncta were unevenly distributed along the dendritic shaft and on the spine necks (Figure 3F).

### H-Link can birth date ECM *in vivo*

To birthdate the ECM *in vivo,* we employed a dual injection paradigm where mouse pups were injected with either the pSYN (Figure 5A) or pCAG driven H-Link AAVs (Figure 5B,C) on P1. HTL549 (1^st^ HTL) was injected into the hippocampus on P6 and HTL646 (2^nd^ HTL) was applied post-fixation. Previous studies report that HTL is cleared from the brain within an hour if there is no HaloTag to bind to [47, 55], thus we can date the ECM to the day of injection. We observed HTL549 signal diffusely throughout the hippocampus with aggregation around certain cells that were infected in both P12 and P16 tissue (Figure 5). Thus, the stability of the probe observed in culture is preserved *in vivo*. We again observed two distinct types of structures: net-like structures (Figure 5B) and coat-like structures (Figure 5C). We observed strong HTL549 signal at proximal processes that appears to be the axon initial segment (Figure 5A’,A’’,C, yellow arrows). We found neuronal soma that were organized early but did not continue to add ECM (Figure 5A’, white arrow), and others that organized early and did continue to add ECM (Figure 5A’’, white arrow and B). We noticed that the HTL549 signal appeared more diffuse than the HTL646 signal (Figure 5C, grey arrows), implying older ECM tends to diffuse away from the neuronal soma. Finally, we noticed distal dendrites tended to have more 2^nd^ HTL than 1^st^ HTL (Figure 5A’,A’’, blue arrows), emphasizing that the progression of ECM clustering toward distal dendrites is a temporally regulated organizational process.

**Figure 5:**
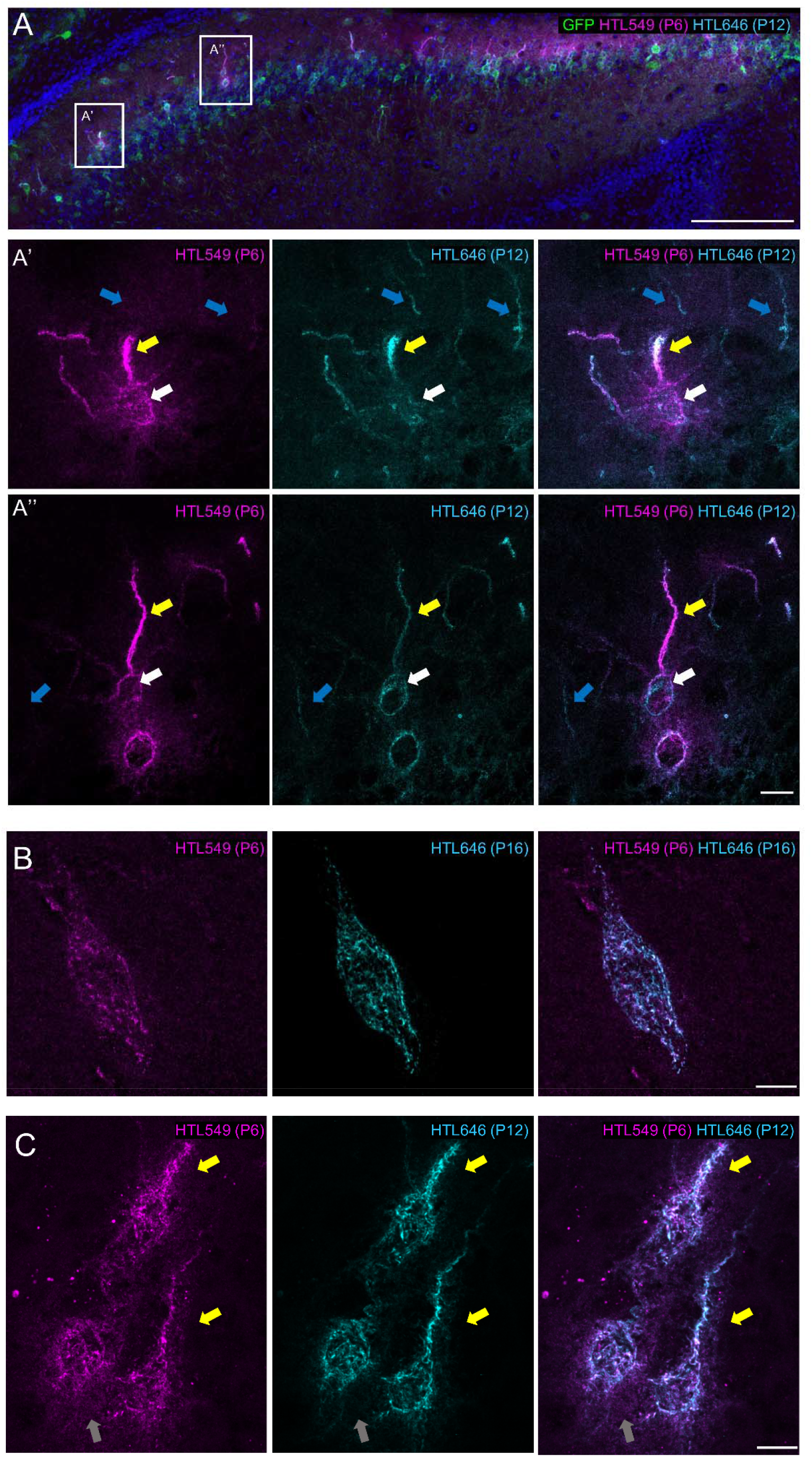
H-Link marked ECM is stable and shows developmental regulation *in vivo*. **(A)** pSYN-eGFP-2A-H-Link AAV injected at P1 followed by HTL549 (1^st^ HTL) hippocampal injection at P6 and sacrifice and post-fixation HTL646 (2^nd^ HTL) staining at P12. The first HTL binds to early produced ECM and is stable for 6 days. The 2^nd^ HTL binds to any unbound H-Link in the tissue. The 1^st^ HTL signal is around infected neurons in the CA1 and fasciola cineria. **(A’)** Inset highlights a neuron which has the 1^st^ HTL near the soma that does not overlap with the 2^nd^ HTL (white arrow), showing stable and early organized ECM. **(A’’)** Inset shows a neuron where soma organization is not completed at P6 and thus there is 2^nd^ HTL segregated from the 1^st^ HTL (white arrow). The brightest 1^st^ HTL structure was the axon initial segment (yellow arrow) in both insets. Scale bar **(A)** 100 µm, **(A’,A’’)** 10µm. **(B)** pCAG-mGFP-2A-H-Link AAV injected at P1 followed by 100 µM HTL549 (1^st^ HTL) hippocampal injection at P6 and sacrifice and post-fixation HTL646 (2^nd^ HTL) staining at P16 showing a conventional net structure of the PNN and the 2^nd^ HTL fills the gaps of the first HTL. **(C)** pCAG-eGFP-2A-H-Link AAV injected at P1 followed by 100 µM HTL549 (1^st^ HTL) hippocampal injection at P6 and sacrifice and post-fixation HTL646 (2^nd^ HTL) staining at P12 shows a coat-like structure with diffusion of the 1^st^ HTL from the 2^nd^ HTL (white arrow) while yellow arrows highlight the axon hillock.

## Discussion

The ECM is consistently implicated in learning and memory, but the physiology has been difficult to understand. While WFA staining displays distinct ECM clusters around a subset of neurons, whether ECM clustering is restricted to WFA (+) neurons is unclear since WFA labels specific CS-GAG [30, 34, 56]. Alternatively, global staining with antibodies against CSPGs or HABP shows less distinct clusters around some cells [23, 57]. However, due to the high level of expression of these ECM components both in the neuronal ECM clusters and interstitial matrix and the dense organization of the brain’s cells, it is difficult to reveal the ECM architecture on individual neurons. Additionally, investigations into ECM organizing principles in culture are hampered by poor staining with WFA until DIV20 and above [15, 22, 48], indicating the slow maturation process of WFA-labeled CS-GAG in cultured neurons. In this study, we targeted HAPLN1, an ECM protein that stabilizes the higher-order HA-CSPG complex, to create H-Link. H-Link allows us to probe the maturation process of ECM organization in culture, sparsely label and visualize ECM structure on individual neurons in the brain, monitor ECM development on cultured neurons, and birthdate the process of ECM deposition by using distinct fluorescent conjugated HaloTag ligands. We saw that H-Link aggregates on neurons, with dense staining in the soma and punctate staining on distal processes, reminiscent of conventional WFA (+) PNNs in the brain. Application of exogenous H-Link to neurons revealed heterogeneity of neuronal coverage, synaptic occlusion of the ECM, aggregation on neurite contact sites, and temporal and spatial regulation of the development of ECM aggregates. Excitingly, we can visualize and follow ECM deposition at much earlier time points than other studies. *In vivo*, we found H-link labels conventional WFA (+) PNNs and novel PNN-like clusters on the hippocampal excitatory neurons. We found regional differences in the ECM and can birthdate the developing ECM *in vivo*.

While H-Link deposition is observed in a larger neuronal population than WFA (+) neurons, H-Link is not assembled on every neuron *in vivo* and *in vitro*. Why is the ECM clustered around some neurons as opposed to other neurons? This is a fundamental question that H-link may answer by revealing organizing principles. For example, the heterogeneity of exogenous H-Link on cultured neurons is quite striking. After analyzing the identity of the neurons, we found clear population-level differences, wherein the hippocampal excitatory neurons had a higher total percentage of neurons with H-Link clusters than the other populations, and a subset of inhibitory neurons displayed a high amount of H-Link signal (Figure 2C-E). The time-lapse imaging showed that there is temporal heterogeneity among ECM organizers: early organizing neurons begin to incorporate H-Link into their ECM as early as DIV5, while late organizers aggregate signal after DIV7. Negative neurons intermingled with early and late organizers remain negative throughout the imaging session. This contrast highlights cell-intrinsic mechanisms of ECM deposition. In support of the cell-intrinsic mechanism, we observed the induction of the H-link aggregation after neurite crossover (Figure 3A-D). The specific spatial and temporal patterns in the presence of continued H-Link in the culture media imply that H-Link is binding to assembled ECM rather than catalyzing the assembly.

*In* vivo, we expected that GFP (+) cells would be decorated with H-link as there should be secreted probes around them. However, we observed GFP (+) neurons without H-Link clusters (Figure 4A, S2E). These cells are predominantly located in the cortex. While the heterogeneity among neurons underscores the specificity of our probe, caution should be taken when interpreting the data following the overexpression of the probe. Prior research favors a direct binding and stabilization role for HAPLN1 to the hyaluronic acid matrix [6, 16, 23, 27, 49], however, it is unclear what is necessary to direct HAPLN1 binding to HA scaffold, nor how stable this interaction is *in vivo*. One possibility is that there is an excess of open binding sites for HAPLN1 in structured ECM, and the H-Link signal reflects its architecture. In this case, lack of signal equates to a lack of structured ECM around that neuron. Another possibility is that HAPLN1 binding sites are pre-occupied by endogenous HAPLNs, and our probe cannot outcompete once structured ECM is stabilized. In this case, our probe can identify structured ECM with the least amount of endogenous HAPLNs to outcompete, and the absence of signal does not equate to the lack of structured ECM. Our culture data showed that the deposition of exogenous H-link on neurons stably increases over a 12-day time course, supporting that H-Link shows the assembly of ECM clusters instead of a dissociation of endogenous link proteins from the structured ECM. AAV-mediated direct expression of H-Link or application of exogenous H-Link exhibited a similar pattern in culture (Figure 1, 2). The H-Link pattern strikingly overlapped with PNN architecture when expressed in WFA (+) neurons *in vivo* (Figure 4A). Moreover, we observed that there are uninfected neurons that can capture H-Link *in vivo* (Figure S2E-G). While the identity of these specialized ECM-organizing neurons is unknown, these observations support that H-Link marks the architecture of the structured ECM. However, we cannot fully exclude the 2^nd^ possibility, and the competition of the probe with endogenous link proteins should be considered when quantifying H-Link signal. Genetic knock-in of the tag into the endogenous locus of link proteins will rule out any potential overexpression artifacts. Taken together, our *in vivo* and culture observations point to molecular organizers of the ECM that are heterogeneously expressed in neuronal populations. Although there is known heterogeneity in the expression level of individual ECM components between regions and cellular subtypes of the brain [58], we have provided an assay intrinsically suited to screen through these possible ECM organizing candidates.

Another feature of H-Link is the ability to use distinct fluorescently conjugated ligands to date when the ECM was produced. In culture (Figure 1C), upon adding HTL646 (2^nd^ HTL) 48 hours after HTL549 (1^st^ HTL), we could find areas where there was 2^nd^ HTL but no 1^st^ HTL, showing new accumulation as expected from the continuous expression of the probe. However, there were areas where neither HTL was observed, adding further evidence for the intrinsic spatial regulatory mechanism of ECM development. A fair amount of overlap of the two signals was observed, which could be attributed to an abundance of binding sites for H-link in a location or an increase in the binding sites over time. The observation of increasing 2^nd^ HTL over time supports the latter over the former. *In vivo*, we showed that HTL549 (1^st^ HTL) injected at P6 can last till P16, again highlighting the stability of developing ECM (Figure 5). By comparing the signal between injected 1^st^ HTL and post-fixation stained HTL646 (2^nd^ HTL), a few patterns emerged. We consistently observed that one of earliest-organized and longest-lasting ECM structures is around the axon hillock, determined by the known stereotyped morphology of hippocampal neurons [59]. Like the culture, there are areas where both HTL signals overlapped, areas where there was signal from either the 1^st^ or 2^nd^ HTL, and areas with no HTL signal *in vivo*. The spatial specificity demonstrates a regulated process of ECM deposition wherein the later ECM fills gaps in the original ECM. Also, we can find different levels of HTL549 on adjacent infected neurons with similar levels of HTL646, showing early organizers and late organizers *in vivo*. This type of birthdating assay will be useful for observing the regulatory process of ECM in conjunction with a behavioral paradigm.

One of the most striking *in vivo* observations is the two distinct ECM structures we found: a net-like structure that is observed in conventional WFA (+) PNNs, and an unwoven fiber coat mostly found around cells in the pyramidal layer of the hippocampus that we have termed a perineuronal coat (PNC). When both types of coating are seen together, the structural differences are obvious (Movie 6). This sharp contrast between types of ECM coating demonstrates the power of our sparse labeling approach. Staining densely packed neurons in the cell body layer with coats could look like an even, diffuse staining pattern, whereas sparse labeling allows us to visualize the aggregation around single neurons. The unique patterns among the hippocampal excitatory neurons suggest that it is unlikely due to random aggregation of the probe. Although H-Link overexpression does not change WFA (+) PNN architecture and CS-56 signal (Figure 4A and S2D), we cannot exclude the possibility that overexpression of H-Link may impact the endogenous architecture of PNCs, which needs further validation. If true, excitatory neuron PNCs challenge the parvalbumin neuron PNN centric model of how the ECM impacts plasticity, and the specific role of each type of ECM would need to be determined.

What is the role of the interstitial matrix in regulating neuronal plasticity? Conventional ECM staining displays a uniform pattern in the interstitial matrix where the neurites reside, and the intestinal matrix is considered a diffuse, unorganized gel-like architecture [6, 14, 15, 18, 60]. In our study, we observed that H-Link clusters are organized along dendrites, which are separated from synaptic proteins and are occluded from the spine heads. Furthermore, we observed an increase in the density and intensity of these dendritic puncta in the hippocampus from 3 weeks to 8 weeks of age, coinciding with developmental maturation and increasing synaptic density [59]. This raises a puzzling question about the function of these puncta. Is the ECM a barrier to synaptogenesis along dendrites? Does the ECM stabilize existing synaptic wiring? Does the process of synaptic development clear the local ECM? Is the ECM fully occluded from the synaptic cleft? Ultrastructural studies are necessary, and by replacing the HaloTag in H-Link with a proximity labeling tool, we could stain the ECM for EM studies.

Taking all our data together, we posit a new model to oppose a conventional view where the impact of the ECM on synaptic plasticity is mostly confined to dense aggregates around PV neurons. Our data reveals ECM around both excitatory and inhibitory somas as well as aggregated ECM at distal dendrites that could play barrier or stabilizer roles. This allows the ECM to play a more dynamic role and may help conceptualize contradictory findings where both digestion and inhibition of endogenous ECM digestive enzymes can cause a similar phenotype, or where digestion can cause opposite plasticity phenotypes in brain regions [5, 47, 61, 62]. Rather than affecting one functional unit, these manipulations affect a balance of ECM factors to cause a phenotype. This model opens a conceptual framework for cell-type-specific and region-specific ECM manipulations to identify organizing principles of how the ECM helps regulate plasticity. Overall, H-Link is poised to spearhead this novel exploration through its modular design to interrogate cell type of interest combined with a reporter of interest, orthogonal fluorescent ligands for longitudinal studies, and in culture screening assays for molecular organizers.

## Methods

### Animal Care

The animal care and procedures followed the standard regulations set by the National Institutes of Health and received approval from the Institutional Animal Care and Use Committee (Protocol 00001695). The C57Bl6/J mice were kept in standard housing conditions at the University of Utah, with unrestricted access to food and water, and subjected to a 12:12 light/dark schedule. Sprague Daley rats were kept in standard housing conditions at the University of Utah, with unrestricted access to food and water, and subjected to a 12:12 light/dark schedule.

### Virus production

AAV for *in vivo* injections and cell culture was produced in lab as previously described [63]. Briefly, HEK cells below passage 10 are plated into a 15 cm dish/virus (Fisher 08-772-24) in DMEM-H (DMEM-h (invitrogen 10569010) + NEAA (Invitrogen 11140050) + 10% FBS (invitrogen 16140071) to achieve 90% confluency the following day. Before transfection, media is changed to DMEM-H with 5% FBS. Cells are transfected with AAV vector: AAV capsid (2/1 or PhP.eB): helper plasmid in a 1:4:2 ratio by weight for a total of 40 µg mixed with PEI (40K MW #24765 Polysciences) in DPBS (Invitrogen 14190144). Media was changed to DMEM-H + 10% FBS and collected at consecutive 72-hour time points, following which cells were scraped (Fisher 08-100-241) from the plate and media and cells were spun at 2000 x g for 15 minutes at room temperature (RT). Virus from cell pellet and media were isolated separately. Media was incubated at 4 °C for 2 hours in 20% PEG solution (40% w/v stock: 2.5 M NaCl, 40% PEG (Sigma P2139) followed by a 20 min 3,000 x g spin at 4 °C. Pellet was resuspended in salt activated nuclease (salt activated nuclease, Articzymes #70910-202) buffer (40 mM Tris-Cl, 500 mM NaCl, 10 mM MgCl_2_, pH 9.5, 25 U/ul SAN) and kept at 4 °C for up to 1 week. Cell pellet was resuspended in SAN buffer, incubated at 37 °C for 1 hour and added to resuspended media pellet and incubated for an additional 30 min and spun at 2000 x g for 10 min at RT. Supernatant was carefully layered onto a prepared iodixanol gradient (15% iodixanol diluted in 1 M NaCl/PBS-MK buffer, 25% iodixanol diluted in PBS-MK buffer with phenol red, 40% iodixanol diluted in PBS-MK buffer, 60% iodixanol with phenol red) (60% Stock Sigma D1556) and spun at 250,000 x g for 3 hours at 4 °C and the 40% fraction was collected. Virus was concentrated to a volume of 0.25 mL in a 100KD cutoff Amicon centrifugal filter (Millipore UFC910096) pretreated with a series of DPBS+ Pluronic F68 (Fisher 24-040-032) washes (0.1% 15 minutes and dump, then 0.01%, 0.001% Pluronic in DPBS, 1000 x g spins for 5 min between each). To determine titer, the virus was treated with DNAse and then a RT-PCR was run. Virus was diluted to 1e^12^ GU/ml in autoclaved filter sterilized PBS for all applications. PHP.eB serotype was used for pCAG H-Link injections and viral transduction of neuronal cultures, for all other applications, AAV 2/1 serotype was used.

### HaloTag ligand and Hyaluronidase preparation

Janelia fluorophore 549 HaloTag ligand (HTL549) was a gift from Erik Jorgensen, University of Utah, at 2 mM stock in DMSO kept at -20C. HTL549 was diluted in water to a concentration of 100 uM for *in vivo* injection and a 10 uM stock for culture and ICC/IHC. Janelia fluorophore 646 HaloTag ligand (HTL646 5ug promega GA1120) was prepared according to manufacturer’s instructions. HTL646 was resuspended in DMSO at 2 mM, and then diluted in water to a 10 µM stock for culture and IHC. Both HTLs were added at a final concentration of 10 nM to neuronal cultures for viral H-Link or exogenous H-Link assays. Hyaluronidase (Sigma H3506) was prepared fresh by diluting 1 mg (1000 U/mg) of powder into 100 µL of neuronal media for a stock solution of 10,000 U/mL and a final concentration of 100 U/mL.

### Exogenous application of H-link probe

HEK293T cells were plated into 12 well plates in DMEM + 10% FBS to reach a confluency of 75% the following day and were transfected using lipofectamine (Fisher 11668027) and 0.5 ug of DNA (vectors: H-link and H-link mutants). The following day, the media was replaced with neuronal maintenance media, and the supernatant was harvested 24 hours later, spun at 200 x g to remove debris, and diluted with fresh media. Media was either incubated with HTL549 (final concentration of 10 nM/well) fresh and then added to neuronal cultures (∼100 uL/well), or aliquoted and frozen. Frozen aliquots were thawed on ice, warmed to 37 °C, incubated with HTL, and added to neuronal cultures. Each batch of HEK293T conditioned media was tested for protein expression via Western Blot against the HaloTag.

### SDS PAGE Western Blot

All protein samples were treated with Gel Loading Buffer (GLB, 25% Glycerol (v/v), 2% Sodium dodecyl sulfate (w/v), 0.5 mg/ml Bromophenol Blue, 5% 2-Mercaptoethanol, 60 mM Tris-HCl, pH 6.8) before running SDS-PAGE. Media and lysate from HEK293T cells was loaded in SDS-PAGE gel in electrophoresis buffer (2.5 mM Tris-base, 19.2 mM Glycine and 0.1% SDS) at 140 V for 1 h. Proteins were transferred to a PVDF membrane in transfer buffer (2.5 mM Tris-base, 19.2 mM Glycine and 10% MeOH) at 100 V for 1 hour. The membrane was blocked in TBST (100 mM Tris-HCl, 150 mM NaCl, 0.1% Tween-20, pH 7.5) containing 5% skim milk for 1 hour and incubated at 4 °C overnight with mouse anti-Halo (1:1000 promega G921A). After washing with TBST, the membranes were incubated with peroxidase conjugated donkey anti mouse (1:10K, Jackson 715-035-151) at RT for 1 hour. For exposure, the membrane was rinsed with TBST and was subjected to SuperSignal TM West Pico PLUS Chemiluminescent Substrate (Thermo Fisher Scientific, PI34580) and analyzed on a ChemiDocTM XRS+ system (Bio-Rad).

### Neuronal Culture

Cortical or Hippocampal cultures were isolated from P0 mouse or rat pups, respectively using standard methods [64]. Briefly, forebrain or hippocampi were dissected in dissection media (1x HBSS,(Gibco 14185-052), Pen/Strep (Gibco 15140-122), 2 mM sodium pyruvate (Gibco 11360-070), 10 mM HEPES (Gibco 15630-080), 30mM glucose) and enzymatically (165U of papain (Worthington LS003119) in 1mM HEPES (Gibco 15630-080), 80 mM Na_2_SO_4_, 30 mM K_2_SO_4_, 5 mM MgCl_2_, 20 mM glucose), and mechanically digested to achieve a single cell suspension. Cells were counted and plated at a density of 100k-300k in plating media (MEM (Gibco 11090-099), 10% Horse Serum(Gibco 26050-088), 20 mM glucose, pen/strep, 1 mM sodium pyruvate, 2 mM Glutamax (gibco 35050-061)) onto prepared 12 or 24 well plates coated with 60 mg/ml Poly D lysine (Millipore A-003-E)) in borate buffer (50mM boric acid, 12.5mM sodium tetraborate in water pH 8.5). For live time-lapse imaging, neurons were plated into glass bottom plates (Fisher 662892)), whereas for ICC neurons were plated onto pretreated coverslips (18 mm coverslips (Fisher cat# 12-545-84 18CIR-ID) incubated in ethanol at RT overnight, washed and dried at 225 °C overnight). After 2-3 hours, plating media was exchanged for neuronal maintenance media (Neurobasal A (Invitrogen cat#10888-022), 0.5 mM Glutamax, 10mM glucose, 2% B27 (Invitrogen 17504-044). Half media changes of maintenance media were performed every 3-4 days to maintain culture health. To express H-Link in culture, AAV was added between DIV1-3 to neuronal cultures. Cultures were checked daily for GFP expression, and upon visible GFP expression (DIV7-9), HTL549 was added, and plates were moved to an incubation chamber for live imaging. HTL646 was added 48 hours after HT549. Exogenous H-Link (unmutated H-Link, untransfected CM and mutant CM) with HTL549 was added on DIV13 and neurons were processed for ICC on DIV14. For hyaluronidase treatment, neurons were incubated with 100 U/mL hyaluronidase for 2 hours before adding H-Link CM. For live-imaging, H-Link CM was added between DIV2-5 and hyaluronidase was added on DIV7.

### Live Imaging

All live imaging was performed on the Keyence BZ-X800 all in one fluorescent microscope with BZ-X Viewer software. For viral overexpression, multi-point z-stacks were acquired every 2.5 hours using a 40x objective and filter cubes: Chroma C211830 for GFP, Chroma 211831 (Cy3) for HTL549, and Chroma C211829 (Cy5) for HTL646 with no binning. Single plane images at each time point were selected manually for best focus and combined into stacks for time-lapse videos. For exogenous H-Link, imaging began within a few hours of adding the exogenous probe. Multi-point stacks were acquired every half hour using a 40x objective and the Cy3 filter cube and white light (BF) for up to two 24-hour imaging sessions or multipoint stacks were acquired every hour using a 20x objective with 2x2 binning for 8 imaging sessions spaced 1 hour apart lasting a total of 12 days. Half media changes were performed in between imaging sessions on DIV3, DIV7 and DIV10. Higher resolution (40x) images were manually compiled for time-lapse videos. For 12-day time lapse, best focus images for each imaging session in each channel were compiled into a time-lapse movie by the BZ-X analyzer software package. For hyaluronidase treatment, the culture was removed on DIV7 from the incubation chamber after completion of the imaging session, hyaluronidase was added at a final concentration of 100 U/mL, and imaging was restarted.

### Immunocytochemistry

Neurons were washed 3x in PBS and then fixed in 4% PFA (EMS 15710), 4% sucrose in PBS at RT. After a 3x PBS wash, cells were permeabilized in 0.02% Triton X (Sigma X-100) in PBS (PBST) at RT followed by blocking solution (PBST + 4% BSA (Sigma A9647)). Neurons were incubated in primary antibody solution (PBST 2% BSA) overnight at 4 °C. Primary antibodies used were chicken anti-MAP2 (1:2000 Abcam ab5392), rabbit anti-neurogranin (1:200 Millipore AB5620) mouse anti-GAD67 (1:100 Millipore 10888-022), guinea pig anti-HOMER1A (1:100 Synaptic Systems 160004), guinea pig anti-VGLUT (1:2000 Millipore AB5905) rabbit anti-SAP102 (1:200 ThermoFisher PA5-29116). After three 5-minute PBS washes, cells were incubated in secondary antibody solution (PBST 2% BSA) at RT. Secondary antibodies used were goat anti-mouse cy5 (1:500 invitrogen A10524), donkey anti-mouse Alexa 488 (1:1000 Jackson Immuno Research) donkey anti-rabbit Alexa 488 or Alexa 647 (1:500 for both, Invitrogen A212026 and Jackson immune Research 711-605-152) goat anti-chicken Alexa 488 (1:1000 Invitrogen A11039), Donkey anti-guinea pig Alexa 488 (1:1000 Jackson Immuno Research 706-545-148) and Hoescht 33343 (1:10K Invitrogen H3570). Following another three 5-minute PBS washes, coverslips were mounted onto slides (Fisher 12-544-2) using Fluoromount gold (Southern Biotech 011-01) and dried overnight before storage at -20 °C.

### Injection into neonatal pups

Injections of virus or HTL549 occurred on P1-P6 as approved by the Institutional Animal Care and Use Committee (Protocol 00001695). Pups were anesthetized on ice for 2-3 minutes, paw squeeze was used to confirm anesthesia. Pups were placed on a metal surface with ice cold water underneath and injected by hand using a 10 µL Hamiltion Syringe (#1701) with a 32 gauge needle (Hamilton 7803-04). Pups were injected as parallel to the skull as possible 2 mm above the eyes along the midline from bregma and 2 mm diagonal from bregma above the eye perpendicular to the skull. The needle was inserted 2-3 mm in, 1.5 µL of virus and/or HTL549 was injected over 30 seconds, and after a 30 second count, the needle was pushed in slightly more for 30 seconds and removed over a 30 second count. Virus was diluted to a titer of 1e^12^ GU/ml in autoclaved filter sterilized PBS, and HTL549 and HTL646 were at a concentration of 100 µM (see HTL preparation). After both injections, pups were placed on a warmed animal heating pad (K&H Manufacturing model # 1060) for at least 10 minutes before being returned to their home cage.

### Perfusion and sectioning

Mice were anesthetized using isoflurane (VetOne 502017) and then transcardially perfused with ice-cold PBS and then ice-cold 4% PFA (Sigma 441244) in PBS. Brains were dissected and post fixed in 4% PFA in PBS overnight at 4 °C. Brains were cryoprotected at 4 °C in 15% sucrose in PBS till they sank, and then moved to 30% sucrose for 48 hours at 4 °C. Brains were imbedded in OCT compound (Ted Pella 27050) and frozen in a mixture of dry ice and ethanol in embedding molds (Polysciences 18646A-1) and left at -80 °C for up to one year. Brains were sectioned coronally at 30 µm using a cryostat () through the hippocampal formation and sections were collected in 50% glycerol in 0.5x PBS roughly 360 µm apart. Samples were stored at -20 °C.

### Immunohistochemistry

Coronal sections were washed three times for 10 minutes in PBST. For *wisteria floribunda agglutinin* (WFA) staining, samples were first blocked using a biotin blocking kit (Vector Labs SP2002). Sections were then incubated in block (PBST +5% Normal Goat Serum (NGS)(Thermo PCN5000), +5% Normal Donkey Serum (NDS)(Sigma D9663)) for 1 hour and then incubated in primary antibody solution(PBST + 2.5% NGS and 2.5% NDS) overnight at 4 °C. Primary antibodies used were chicken anti-GFP (1:500 Abcam ab13970), mouse anti-CS-56 (1:200 Sigma C8035), biotinylated WFA (1:1000 Vector Labs B-1355). Sections were washed three times for 10 minutes in PBS and then incubated for 1 hour at RT in secondary antibody solution (0.5x blockPBST +2.5% NGS and 2.5% NDS). Secondary antibodies used were goat anti-chicken Alexa 488 (1:1000 Invitrogen A11039), 10 µM HTL646 (1:500), Streptavidin-cy3 (1:1000 Jackson Immuno Research 016-160-084) and Hoescht 33343 (1:10K Invitrogen H3570). sections were washed three times for 10 minutes in PBS, and then placed on slides. Sections were mounted with flouromount gold and coverslips (TedPella 260152) and left to dry overnight at RT and stored at -20 °C.

### Imaging

Immunocytochemistry and immunohistochemistry images were captured either on a Leica SP8 confocal microscope with LASX software, Zeiss 700 confocal microscope or a Zeiss 880 Airyscan confocal microscope with Zen Black software at the Fluorescence Microscopy Core Facility located at the University of Utah. All images were subjected to processing and analysis utilizing the open-source software Fiji, accessible at.

### Quantification and Statistical analysis

All images were analyzed in the open-source software Fiji, and the variations in all results were determined using GraphPad Prism9 software.

### Quantification of neuronal H-Link coverage in excitatory or inhibitory neurons in hippocampal or cortical cultures

All images for quantification were taken at the same image settings and magnification. 4 channel max projection images were used. Regions of interest (ROIs) were drawn close to the outer border of each neuron as determined by the excitatory or inhibitory marker signal blinded to the H-Link signal. ROIs were categorized as either inhibitory or excitatory based on neurogranin or GAD67 signal. Neurons that were challenging to categorize were excluded from the quantification. ROIs were measured for mean intensity in the H-Link channel. Mean intensities were graphed as a cumulative distribution for each culture type. For statistical testing, a Kolmogorov-Smirnov test was performed on the between neuron types in different cultures to compare cumulative distributions. For % of neurons, ROIs were drawn around neurons blinded to the H-Link channel, categorized, and then, in the H-Link channel, manually counted as positive or negative. Counts for each culture were converted to percentages and used for further statistics and graphing. Two-way ANOVA with culture type factor and neuron type factor was used. ROIs for cumulative distribution were drawn blinded to culture type, whereas manual counts were done unblinded to culture type and done by different experimenters. The same images were used for both quantifications. 3 cultures per condition (cortical and hippocampal) 3 images per coverslips, 10 coverslips (Blinded: 386 excitatory neurons and 220 inhibitory neurons unblinded: 659 excitatory neurons, 420 inhibitory neurons) in the cortical cultures, 9 coverslips (Blinded: 534 excitatory neurons and 326 inhibitory neurons, unblinded: 2001 excitatory neurons, 396 inhibitory neurons) for hippocampal cultures.

### Quantification of total H-Link signal in H-Link CM vs mutant H-Link CM, CM, HTL only and hyaluronidase pretreatment

All images were taken with the same image settings and magnification. Single plane images in the H-Link channel were measured for total fluorescence intensity. 3 images per condition per culture were taken (6 total images). Each image intensity was normalized to the average H-Link intensity of the same culture. A repeated measures ANOVA as used as data passed normalcy test (Kolmogrov-Smirnov test)

### Quantifying neuronal H-Link signal over 12 day time-course

Stacks from each imaging session were concatenated and ROIs were drawn to encompass each neuron throughout the time-course. If the neuron moved significantly, a second ROI was drawn to accommodate the movement, and so forth for continued movement. ROIs were measured using the multi-measure command in the ROI manager for mean H-Link signal at every time point. The background was calculated by drawing small ROIs around seven areas of low signal and measuring the mean at every time point. The minimum value from the seven background ROIs was subtracted from the mean signal of each neuronal ROI to calculate the corrected mean fluorescence intensity (CMFI). Among 6 fields of view analyzed, all had negative, early and late ECM organizers, thus the field of view shown is representative.

### Quantifying loss of H-Link signal after hyaluronidase signal

20 neurons were chosen that had signal before hyaluronidase treatment. ROIs were drawn close to the neuronal membrane and measured 1 hour before and 7 hours after hyaluronidase treatment for mean H-Link signal. Background was calculated by drawing an ROI near the neuron of interest at the corresponding timepoint and subtracting the measured mean to obtain CMFI. Paired Student’s t-test was used with unequal variance.

### Neuronal H-Link dendritic cluster

All images were taken with the same image settings. Quantification was not blinded. Eight to ten neurons showing the intact dendritic branches in the section (GFP) and high H-Link signal in the cell body were chosen and averaged per animal. Four animals per group for a total of 36 neurons (3 weeks) and 35 neurons (8 weeks) were used. Line ROIs with a width of 3 pixels were drawn along neurites using both H-Link and GFP signal and intensity was measured along the line at the same intervals for all neurons. Background was determined for individual neurons by drawing a smaller line ROI with a width of 3 near the neuron in an area of background signal. The corrected fluorescence intensity was obtained after subtracting the average background signal. The signal was normalized to the highest signal around the soma, and the highest signal was set to the origin. A Two-way repeated measure ANOVA with distance factor and age factor was used.

### Cloning

Mouse Hapln1 (NM_013500) was cloned into the pAAV vector by PCR with the following primers and ligase (NEB) or In-Fusion cloning (Takara). The sequence of the constructs was confirmed by Sanger sequencing.

The final H-Link construct contains LeftITR…*MCS*-CAG2/Syn promoter-*FseI*-mGFP/eGFP-*BsrGI*-T2A-SP-*EcoRI*-HaloTag-*HpaI*-Gly linker-matHapln1-*AscI*-WPRE-poly(A)…RightITR. The restriction enzyme sites are italicized.

- SP: endogenous signal peptide of Hapln1 (MRSLLLLVLISVCWAD, NM_013500)
- T2A: GSGEGRGSLLTCGDVEENPGP
- Gly linker: GGGGG
- matHapln1: mature Hapln1 (NM_013500)
- HaloTag: GenBank: HM157289
- mGFP: membrane (MARCKS mem tag, NM_002356.7) GFP
- eGFP: EGFP (pAcGFP1-N1, Clontech)

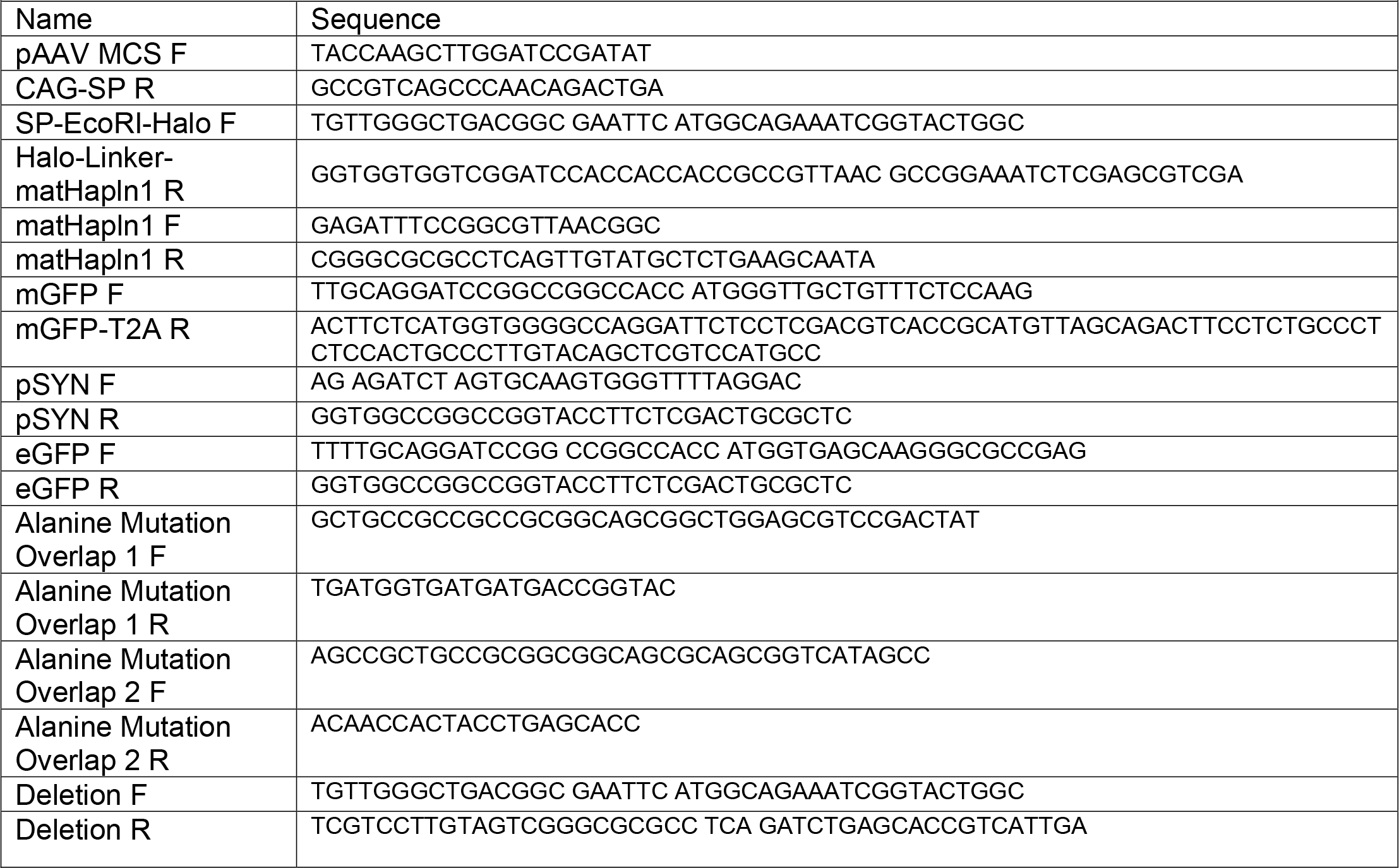

Vectors used in paper

- pAAV-pCAG-mGFP-T2A-SP-HALO-HAPLN1 (pCAG-mGFP-2A-H-Link)
- pAAV-pCAG eGFP T2A SP HALO HAPLN1 (pCAG-eGFP-2A-H-Link)
- pAAV pCAG mGFP T2A SP HALO HAPLN1_del68AA truncation (pCAG-mGFP-2A-H-Linkdeletion)
- pAAV pCAG mGFP T2A HALO HAPLN1-Z616-622AA mutation (pCAG-mGFP-2A-H-Linkalanine)
- pAAV pSYN eGFP T2A SP HALO HAPLN1 (pSYN-eGFP-2A-H-Link)
- pAAV pSYN mGFP T2A SP HALO HAPLN1 (pSYN-mGFP-2A-H-Link)
- pAAV pSYN SP eGFP HAPLN1 (pSYN-G-Link)

**Figure S1 Related to Figure 2:**
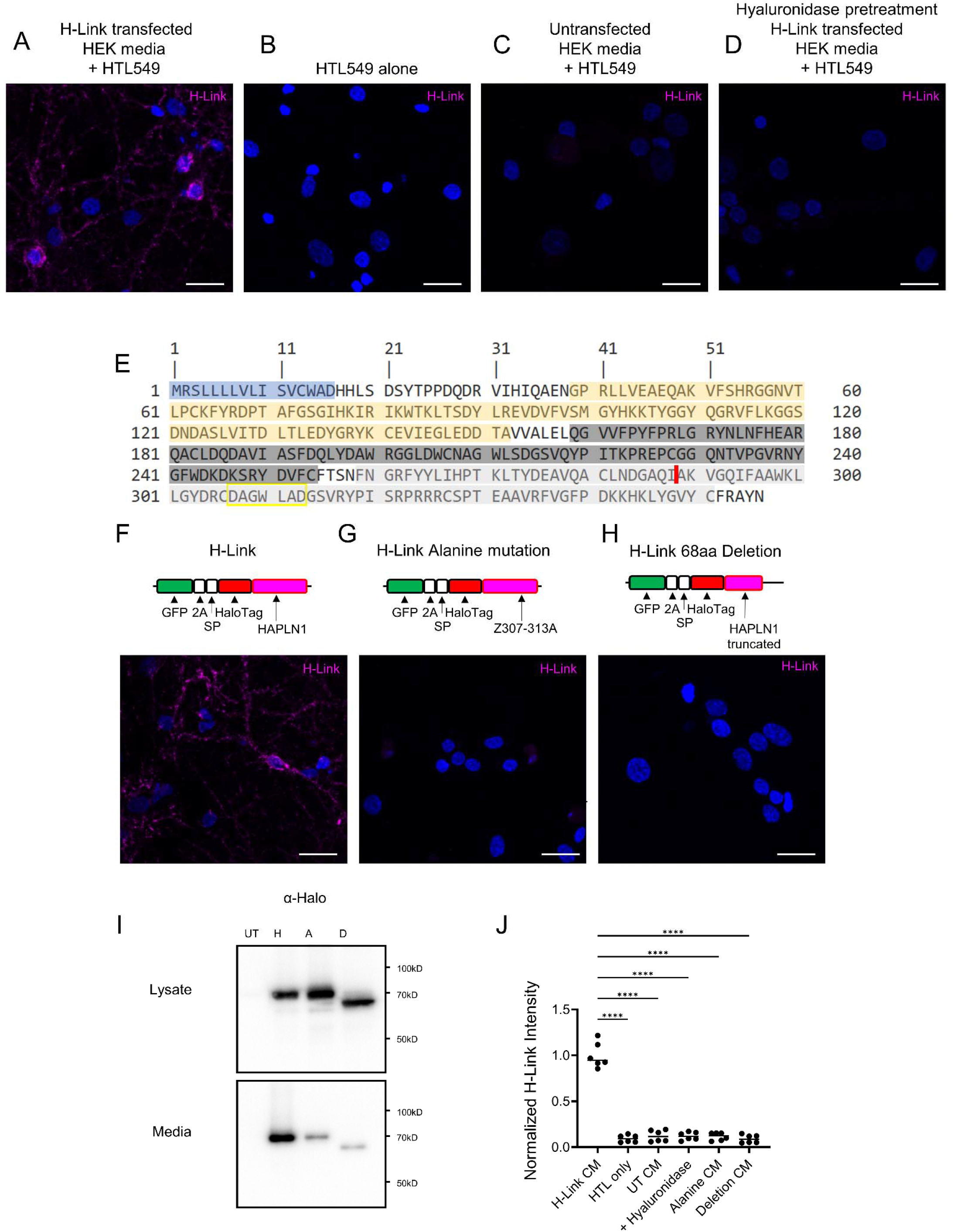
Exogenous H-Link signal is specific and relies on HA binding. **(A-D)** Representative images after overnight incubation with indicated treatment. **(D)** 1 hour hyaluronidase pretreatment at 100 U/mL. Scale bar 10 µm. **(E)** Amino acid sequence of HAPLN1 with domains highlighted: signal peptide (blue), immunoglobulin domain (gold), HA binding domain 1 (dark grey), HA binding domain 2 (light grey). Yellow boxed amino acids are mutated to alanine for H-Link alanine mutation, and a premature stop codon was inserted at the red line for H-Link deletion mutation. **(F-H)** Representative images after overnight incubation with the indicated CM, 5x the amount of CM for mutants. Scale bar 10 µm. **(I)** Western blot of HEK293T cell lysate and media, either untransfected (UT), or transfected with H-Link (H), H-Link with alanine mutation (A), or H-Link with premature stop codon (D) with an anti-Halo antibody shows expression and secretion of all H-Link mutants. The secretion of the H-Link mutants was reduced compared to H-Link. **(J)** Quantification of A-D, G,H in a 2.5 mm area normalized to average total H-Link signal. N=2 cultures, 3 replicates per culture, n=6 images per condition. RM One-way ANOVA with Dunnet’s multiple comparisons. Treatment factor F= 213.4 P<0.0001, all multiple comparisons p<0.0001

**Figure S2 related to Figure 4:**
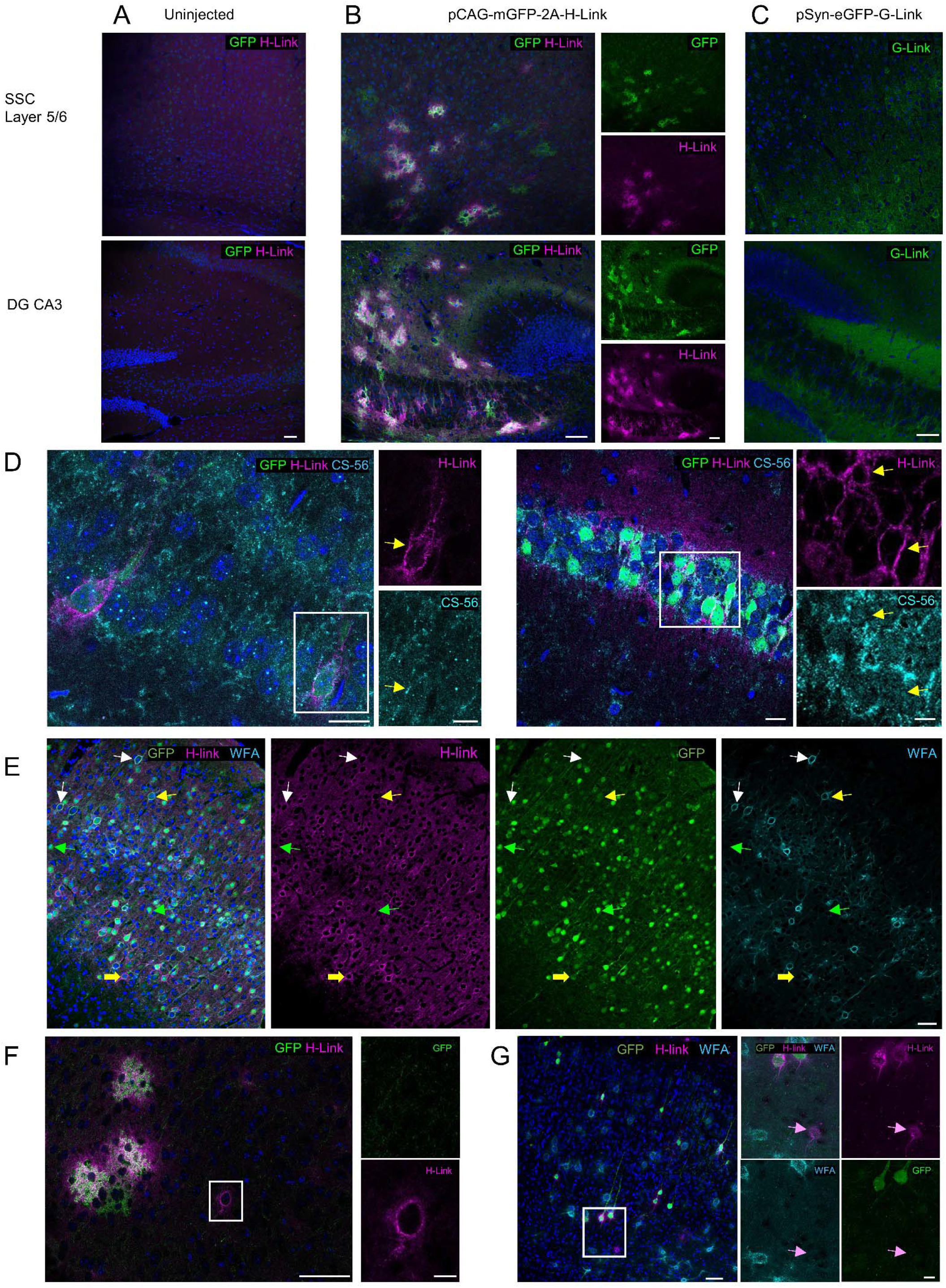
H-Link marks ECM architecture and does not disrupt endogenous ECM in vivo. **(A-C)** Representative images of the somatosensory cortex (SSC) layers 5/6 (*Top*) and the dentate gyrus and CA3 region of the hippocampus (DG CA3) (*Bottom*) of 3-4-week-old mice with no injection **(A)**, injection of pCAGGS (pCAG, a constitutive promoter)-driven H-Link with a membrane GFP reporter **(B)** or a GFP-HAPLN1 fusion, or G-Link driven by the synapsin promoter (pSYN) **(C)**. Staining for H-Link in uninjected tissue shows low background and no specific signal whereas in the injected tissue **(B)** there is expression of the reporter and aggregation around neurons and astrocytes in both brain regions. *Right* is single channel of *Left*. G-Link **(C)** showed a similar staining pattern to H-Link, highlighting the modularity and versatility of the tool. Scale bar 50 µm **(D)** Representative images of CS-56 staining in the dentate gyrus (DG) (*Left)* and CA1 (*Right*) of the hippocampus in P23 mice injected at P2 with pSYN-eGFP-2A-H-Link AAV shows H-Link aggregation does not cause undue aggregation of other ECM components. *Left:* signal is similarly diffuse around the H-Link positive cells. *Right:* arrows point to areas of H-Link aggregation that do not correspond to CS-56 aggregation. Scale bar 20 µm, inset 10 µm. **(E)** H-Link shows heterogeneous coverage of neurons in the somatosensory cortex. Representative max projection of an 8-week-old somatosensory cortex injected with pSYN-eGFP-2A-H-Link AAV at P2. Yellow arrows show triple positive (H-Link (+), WFA (+), GFP (+) neurons, white arrows show neurons that are WFA (+) H-Link (+) but GFP (-), yellow block arrows show H-Link (+) GFP (+) but WFA (-) and green arrow is GFP only cell. Scale bar 10 µm. **(F)** Image of pCAG-mGFP-2A-H-Link AAV injected at P0 imaged at P20 showing expression in astrocytes in the somatosensory cortex. There is H-Link signal around the infected astrocytes and H-link signal around a neuron 50 microns away. Inset shows neuron that is not infected but is able to recruit diffusible H-Link into its ECM. **(G)** 8-week primary visual area pSYN-eGFP-2A-H-Link AAV injected at P2 with inset of layer 5 neurons that are H-Link+, but one is GFP (-) (arrow), and all 3 are WFA (-) (**F, G)** Scale bar 50 µm, inset 10 µm.

**Movie 1. Related to Figure 1C**

The increase in the HTL646 signal indicates the synthesis and incorporation of the probe. The continued presence of the HTL549 signal highlights the stability of the deposited ECM. AAV Infection: DIV1; HTL549: DIV7; HTL646: DIV9. Time lapse imaging: DIV10-11 every 2.5 hours.

**Movie 2. Related to Figure 3A**

H-Link aggregation is heterogeneous around neurons with neurons 1 and 6 having early aggregation around DIV5, neuron 4 having late aggregation after DIV7 and neuron 8 with no aggregation throughout the 11 day time course, Rat hippocampal culture, H-Link CM: DIV2, *Top*: H-Link Brightfield Merge, *Bottom*: H-Link only.

**Movie 3. Related to Figure 3D**

Example of contact-mediated induction of ECM deposition. Soon after growth cone contacts neurite in the top left, there is aggregation of signal. As well, previous contact sites have signal throughout the imaging session. DIV6, H-Link CM: DIV5.

**Movie 4. Related to Figure 3D**

Example of contact-mediated induction of ECM deposition. Start is 5 hours after H-Link CM addition, arrows point to areas of crossover that aggregate signal afterwards.

**Movie 5. Related to Figure 4A**

Imaris visualization of max projection of Figure 4A (*Bottom*), Magenta H-Link, Cyan WFA

**Movie 6: Related to Figure 4B**

Imaris visualization of max projection of Figure 4B (*Bottom*), Magenta H-Link, Cyan WFA

## Acknowledgment

We are grateful to Dr. Megan Williams (University of Utah) for discussion and assistance. We are grateful to Dr. Erik Jorgensen (University of Utah) for providing a HaloTag ligand and Dr. Veronika Romero (University of Utah) for assistance with neuronal culture and live-cell imaging. This work was supported by the Brain Research Foundation (BRFSG-2022-07) to SP and R01NS102444 to SP.

